# HAPPI GWAS: Holistic Analysis with Pre and Post Integration GWAS

**DOI:** 10.1101/2020.04.07.998690

**Authors:** Marianne L. Slaten, Yen On Chan, Vivek Shrestha, Alexander E. Lipka, Ruthie Angelovici

## Abstract

**Motivation:** Advanced publicly available sequencing data from large populations have enabled informative genome-wide association studies (GWAS) that associate SNPs with phenotypic traits of interest. Many publicly available tools able to perform GWAS have been developed in response to increased demand. However, these tools lack a comprehensive pipeline that includes both pre-GWAS analysis such as outlier removal, data transformation, and calculation of Best Linear Unbiased Predictions (BLUPs) or Best Linear Unbiased Estimates (BLUEs). In addition, post-GWAS analysis such as haploblock analysis and candidate gene identification are lacking.

**Results:** Here, we present HAPPI GWAS, an open-source GWAS tool able to perform pre-GWAS, GWAS, and post-GWAS analysis in an automated pipeline using the command-line interface.

**Availability:** HAPPI GWAS is written in R for any Unix-like operating systems and is available on GitHub (https://github.com/Angelovici-Lab/HAPPI.GWAS.git).

## 1 Introduction

Recent advances and publicly available sequencing data of large populations have enabled informative GWAS. As a result, traits of interest have been linked to specific genomic loci unraveling the genetic architecture of complex traits. Demand to run GWAS on large datasets and user-friendly, flexible platforms is an increasingly important demand to fulfill. However, tools have not emphasized the incorporation of pre-GWAS and post-GWAS analysis in combination with GWAS in a holistic tool with comprehensive summary tables and figures.

A past effort has focused heavily on ease of usability. GWAS programs such as GAPIT (Lipka et al., 2012) incorporate a variety of statistical models into a single R package, while others such as FarmCPU (Liu et al., 2016) implement novel statistical models. Other programs use graphical user interfaces (GUIs) such as TASSEL (Bradbury et al., 2007) or web-based platforms such as GWAPP (Seren et al., 2012) and easyGWAS (Grimm et al., 2017). However, these tools do not provide all crucial steps: pre-GWAS (outlier removal, transformation, and BLUP/BLUE calculations) and post-GWAS (haploblock analysis and gene extraction and identification), in addition to a user-friendly platform.

In response to these needs, we have developed Holistic Analysis with Pre and Post Integration (HAPPI) GWAS which provides a complete GWAS pipeline including pre-GWAS, GWAS, and post-GWAS analysis in a single tool. HAPPI GWAS incorporates pre-GWAS steps most beneficial for the data structure of plant-related traits, but also runs generic analyses such as GWAS and post-GWAS analyses. With properly formatted input and the ability to opt in and out of certain analyses in the pipeline, HAPPI GWAS will run data from any species. A summary of the main contributions of HAPPI GWAS include: 1) eliminating the need for multiple tools by providing a comprehensive GWAS pipeline for all phases of a GWAS analysis, 2) allowing high-throughput analysis of multiple traits with easy comparison of GWAS results across all traits and concise, publication-ready figures and tables, and 3) allowing user-defined models and threshold parameters specified at the start of the workflow and automatically implemented throughout the pipeline without additional configuration.

## 2 Implementation

HAPPI GWAS is implemented in four main steps: pre-GWAS, GWAS, post-GWAS, and Outputs, Summaries and Visualizations (for HAPPI GWAS workflow refer to Figure 1). Each step is customizable by the user through a YAML file. The YAML file instructs HAPPI GWAS of the name and location of data input and output and allows for user-defined parameters at each step of the pipeline. Additional information regarding each step, parameter flexibility, and tutorial datasets can be found in the HAPPI GWAS manual (Supplementary Document S1) and on the wiki page linked in the GitHub repository.

**Fig. 1.**
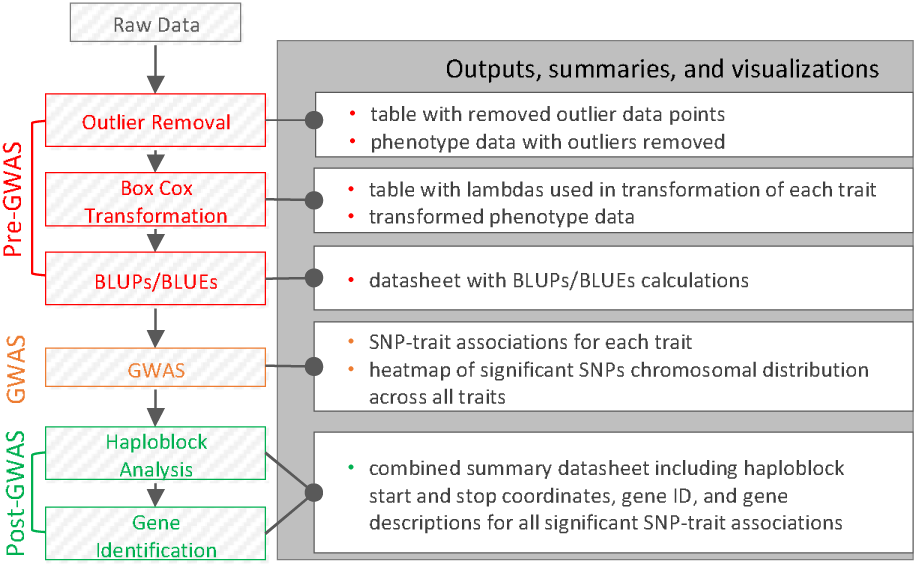
HAPPI GWAS workflow outlining pre-GWAS, GWAS, post-GWAS, and outputs, summaries, and visualizations steps.

### Pre-GWAS

Ensuring raw data meets all assumptions prior to GWAS is vital to reproducible and accurate results but can be difficult to navigate. HAPPI GWAS automatically inputs raw data into pre-GWAS analysis by removing outliers using Studentized deleted residuals (Cook, 1977) and transforming data using the Box-Cox procedure (Box & Cox, 1964). The variance between replicates is evaluated via mixed models to calculate BLUPs and BLUEs, which is particularly important in plant GWAS where several biological replicates are common. Model flexibility allows users to define specific random and fixed effects in these mixed models. If preprocessing is completed externally, the option to skip the pre-GWAS step is available.

### GWAS

The GWAS step accepts user-defined phenotype data and genotype data. Provided genotype files can be used in combination with user-supplied phenotype data. Multiple traits can be stored in each phenotype dataset and run consecutively. Configuration files are edited by the user to supply values for mandatory defined variables that are called when the program is invoked. All GWAS analysis is performed by calling the GAPIT v3 R package (Wang & Zhang, 2018).

### Post-GWAS

Most GWAS packages output a list of significant SNP-trait associations. However, a list of obscure SNP IDs is often uninformative until associated with genes. In the post-GWAS step, a list of significant SNPs is fed directly into a haploblock analysis in Haploview (Barrett et al., 2004). The Haploblock analysis filters SNPs at a 5% minor allele frequency (MAF) and quantifies the degree of linkage disequilibrium (LD) using D prime in the surrounding genomic region to estimate a haploblock (defined as regions of high LD) described with a start and stop location. Genes contained or partially contained within haploblocks are identified and output with respective gene descriptions in the final summary datasheet (Supplementary Table S1-S3). If no genes overlap the haploblock or the significant SNP does not fall within a haploblock, the gene directly upstream and downstream of the significant SNP is given. HAPPI GWAS allows users to define the window size for LD calculations to increase gene identification in a larger interval or to skip the post-GWAS step entirely in species where limited genomic information is available.

### Outputs, Summaries, and Visualizations

GWAS results from all traits found in the phenotype file are summarized concisely in tables and figures as part of the automatic summary output. Collective analysis of related traits can be powerful in the detection of pleiotropy. HAPPI GWAS compiles GWAS results creating a combined GWAS results summary that includes significant SNP IDs, gene names, gene descriptions, and haploblock information (Supplementary Table S1). Two additional summary tables are created: a table summarizing the top five SNP-trait associations with the lowest *P*-value across all analyzed traits (Supplementary Table S2) and a table summarizing the most recurring SNP-trait associations across all analyzed traits (Supplementary Table S3). Lastly, a unique HAPPI GWAS visualization representation of the chromosomal distribution of all significant SNP-trait associations (Supplementary Figure S1) for all traits is provided. This figure is unique to HAPPI GWAS and differs from other multi-trait GWAS visualizations such as Zbrowse (Ziegler et al., 2015) because only significant SNPs are visualized. This novel format allows for easier comparison of genome-wide SNP distributions across the traits as compared to overlapping Manhattan plots. Automatic GAPIT output, as found in the GAPIT3 manual (Wang & Zhang, 2018), is also included in the output.

## 3 Performance Test

HAPPI GWAS excels at multi-trait analysis. Automatic parallelization is implemented to support the excess computational burden that arises from the multi-trait analysis feature but requires a Unix platform. Due to restraints in packages used in parallelization, users wishing to run HAPPI GWAS on a Windows machine must use a virtual machine and install CentOS/Ubuntu (see Supplementary Document S1 for more information).

To further show the computational performance of HAPPI GWAS and dissect the effects of sample size and SNP number on runtime, we tested HAPPI GWAS on a CentOS machine with 500 GB of RAM and 30 TB of disk space. We analyze one trait using filtered genotypic data from the *Arabidopsis* 1001 dataset (Alonso-Blanco et al, 2016) with varying sample size and SNP number with one processing core (Supplementary Table S4). The filtered data has a total of 1,057,383 SNPs and the phenotypic data is from 901 individuals in replicates of two. To run the entire dataset through HAPPI GWAS takes 6 hours and 39 minutes. As the number of individuals in the population decreases, run time remains relatively constant. Using the full genotypic data with 1,057,383 SNPs but decreasing the number of individuals by half (450 individuals) results in a runtime of around 6 hours and 6 minutes. Conversely, decreasing the number of SNPs to half (i.e., to 528,692 SNPs) while maintaining a population size of 901 results in a shorter runtime of 2 hours and 7 minutes.

## 4 Conclusion

HAPPI GWAS is a holistic tool that integrates pre-GWAS, GWAS, post-GWAS, and outputs, summaries, and visualizations. Incorporation of all four steps leads to a comprehensive pipeline that aims to be computationally approachable, regardless of user background. It improves upon past GWAS tools by increasing the scope of analysis and plasticity of defined parameters.

## Acknowledgements

The authors wish to acknowledge Sarah Turner-Hissong for her assistance in early pipeline development.

