## Supplementary Document S1 for "HAPPI GWAS: Holistic Analysis with Pre and Post Integration GWAS"

User Manual for

**HAPPI GWAS**

**H**olistic **A**nalysis with **P**re and **P**ost **I**ntegration **GWAS**

Last updated in May 2020

**Disclaimer:** HAPPI GWAS source code has been extensively debugged and tested. Results should be correct and replicable. However, we do not guarantee certain results for any data. We encourage users to validate results with external tools.

**Supporting Documents:** Source code, demonstration scripts, tutorial data, and support documents including a wiki page, for this package can be found at <https://github.com/Angelovici-Lab/HAPPI.GWAS.git>. Additional *Arabidopsis* reference files can be found on CyVerse (/iplant/home/angelovici_lab/HAPPI_GWAS/).

**Citation:** Multiple statistical methods are implemented. Citation of HAPPI GWAS may vary based on versions used in the analysis. Please refer to citations below:

| **Method** | **Method paper** | **HAPPI GWAS Implementation** |
| --- | --- | --- |
| Studentized Deleted Residuals | Cook, R.D., et al. (1977) Detection of influential observation in linear regression, *Technometrics*, 19.1, 15-18. | Outlier removal |
| Box Cox Transformation | Box, G. E., & Cox, D. R. (1964) An analysis of transformations, *Journal of the Royal Statistical Society: Series B (Methodological)*, 26.2, 211-243. | Phenotype data transformation |
| Mixed Linear Model (MLM) and Population Parameters Previously Determined (P3D) | Yu, Jianming, et al. (2006) A unified mixed-model method for association mapping that accounts for multiple levels of relatedness. *Nature genetics,* 38.2. 203-208. | GWAS |
| Compressed Mixed Linear Model (CMLM) | Zhang, Z., et al. (2010). Mixed linear model approach adapted for genome-wide association studies. *Nature genetics*, *42*.4, 355. | GWAS |
| FarmCPU | Liu, X., et al. (2016) Iterative usage of fixed and random effect models for powerful and efficient genome-wide association studies, *PLoS genetics*, 12.2, e1005767. | GWAS |
| Haploview | Barrett, J.C., et al. (2004) Haploview: analysis and visualization of LD and haplotype maps, *Bioinformatics*, 21.2, 263-265. | Haploblock analysis |

**Contents**

1. INTRODUCTION________________________________________________
   1.1. Why HAPPI GWAS?
   1.2. Getting Started
   1.3. How to use the User Manual
2. DATA______________________________________________________________
   2.1. Phenotypic Data
   2.2. Genotypic Data
        2.2.1. Hapmap format
        2.2.2. Numeric Format
        2.2.3. Naming and Importing Large Genotypic Data
3. ANALYSIS___________________________________________________
   3.1. Pre-GWAS analysis
        3.1.1. Outlier Removal
        3.1.2. Box-Cox Transformation
        3.1.3. BLUPs/BLUEs
               3.1.3.1. BLUPs
               3.1.3.2. BLUEs
   3.2. GWAS
        3.2.1. Models
        3.2.2. Additional GWAS options in GAPIT
   3.3. Post-GWAS
        3.3.1. Haploblock Analysis
        3.3.2. Identify Genes
4. RESULTS__________________________________________________________
   4.1. Intermediate output files
   4.2. GWAS output
        4.2.1. HAPPI GWAS summary figures and tables
        4.2.2. Additional output
5. TUTORIALS_________________________________________________________
   5.1. Editing the YAML file
        5.1.1. Inputting raw data
        5.1.2. Inputting BLUP/BLUE data
        5.1.3. Editing paths in the YAML file
        5.1.4. Filtering by P-*value* or FDR
        5.1.5. Editing the model
               5.1.5.1. Number of variables in the model
               5.1.5.2. Fixed vs random variables in the model
   5.2. Tutorial Datasets
        5.2.1. Maize Demo
        5.2.2. *Demo Arabidopsis* 360 Population
        5.2.3. *Arabidopsis* 1001 Population
6. APPENDIX__________________________________________________________
   6.1. Properties of tutorial files
   6.2. Formatting user input GFF file
   6.3. Formatting user input vcf file
   6.4. Frequently asked questions

REFERENCES

1. **INTRODUCTION**
   1.1. ***Why HAPPI GWAS is important and needed***

Recent advances and publicly available sequencing data of large populations coupled with the development of improved statistical methods has enabled informative genome-wide association studies (GWAS). As a result, the genetic architecture of many traits of interest have been associated with specific genomic loci. Demand to run GWAS, not only on large datasets, but also on a user-friendly, flexible platform has grown and become an increasingly important demand to fulfill.

The increased demand for analyzing large genotypic datasets has been answered with an increase in publicly available tools and methods. However, understanding how to use the plethora of tools for each step (and what steps to perform) in the GWAS analysis can often be a time-intensive, arduous process that requires self-teaching of many tools and extensively formatting output data from one tool to input into the next.

Despite many publicly available tools that conduct GWAS, these tools lack critical steps in the GWAS analysis, such as pre-GWAS (outlier removal, data transformation, and BLUP/BLUE calculation) and post-GWAS analysis (haploblock analysis and candidate gene identification). Additionally, these tools often require that the analysis is run through web-based platforms and workflows that restrict users to set parameters and models. Here, we identify the potential in making an R-based tool that encompasses four steps: pre-GWAS, GWAS, post-GWAS analysis, and outputs, summaries, and visualizations compiled in one tool. Unlike other GWAS tools, HAPPI GWAS provides a comprehensive GWAS pipeline that results in a list of putative candidate genes while maintaining user flexibility throughout the workflow.

1.2. ***Getting started***

HAPPI GWAS is an GWAS tool bundled as an R package that runs on the command-line interface of any Linux and Mac operating system. Due to restraints in packages used in parallelization, users wishing to run HAPPI GWAS on a Windows machine must use a virtual machine and install CentOS/Ubuntu. HAPPI GWAS source code is loaded into the R environment using the R script command in the terminal while outsourcing analysis to external tools (external tools require Python3, ImageMagick, and Java). HAPPI GWAS is free and publicly available for download. Before cloning the HAPPI GWAS GitHub repository, the *devtools* package and the *HAPPI.GWAS* R package must be installed in the R environment (see below). From the terminal, users need to create a file named “HAPPI_GWAS”. Users should take note of the absolute path of this first folder by using the *pwd* command and navigating to that directory. Once in the “HAPPI_GWAS” directory, users can clone the repository from GitHub. Commands are provided as follows:

```
#completed in the R environment:
install.packages("devtools", dependencies = TRUE)	#install devtools package
devtools::install_github("Angelovici-Lab/HAPPI.GWAS")  #install HAPPI.GWAS R package

#completed from the terminal:
mkdir HAPPI_GWAS	   #make initial directory
pwd		   #find absolute path
cd HAPPI_GWAS	   #navigate into directory
git clone https://github.com/Angelovici-Lab/HAPPI.GWAS.git	      #clone repository
cd HAPPI.GWAS	   #navigate into cloned directory
```

When the download process is finished, the next step is to run the setup script (this is only required after the repository is freshly cloned). Output directories are automatically created to store and organize results. To run the setup.R script, type the following command line:

Rscript setup.R

After the setup is complete, users can print the directory structure using *tree* command (if installed on the machine). Printing out the directory structure is not mandatory, but it is a good idea to verify if all the necessary folders are created correctly. The HAPPI GWAS directory structure is shown below:

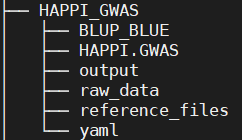

The easiest way to run HAPPI GWAS is to run the tutorial (Demo) YAML files. Here is example code running the maize Demo data. It will run the GLM model using the maize phenotype data provided. (see *Tutorials* 5.1).

Rscript HAPPI-GWAS.R -GAPIT -extractHaplotype -searchGenes Demo_GLM.yaml

1.3. ***How to use the manual***

The next 2-4 chapters will cover input data, options for GWAS analysis (including model and package flexibility), and output results. Chapter 5 presents the maize and *Arabidopsis* tutorial data. Chapter 6 presents frequently asked questions and troubleshooting.

2. **DATA**
   2.1. ***Phenotypic data***

The phenotypic data must be provided by the user in the form of raw data or BLUP/BLUE data. Users are not limited by the number of traits that can be run in HAPPI GWAS. All phenotypes should be saved in a tab-delimited text file (.txt) or comma-separated values file (.csv) with missing data indicated by “NA”. Duplicate values are not allowed in the phenotypic data, so it is necessary to solve duplication problems using mathematical approaches such as *lsmean* or arithmetic mean before feeding the data into the tool. If negative phenotypic values are present in the dataset, the *Family* option in the *powerTransform* function must be changed from “bcPower” to “bcnPower”. This can be done by editing the source code in the func_generate_BLUE.R (line 112) and func_generate_BLUP.R (line 112); both scripts can be found under the HAPPI.GWAS/R directory. (See section 3.1.2 Box-Cox Transformation for additional information).

The raw data and BLUP/BLUE data should have the following format:

*Raw Data*

1. First column: Accession ID/Taxa name should be the first column of the phenotype file.
2. Column 2 - # of variables: these columns can be used to add additional variables in the model.
3. Remaining columns: should contain the observed phenotype measurements from each accession where each column designates one trait. Column names will indicate phenotype names to be used in the remaining analysis.

An example of raw *Arabidopsis* phenotype data is provided below where column 1 is the Accession ID, column 2 is the replicated phenotype measurements (per accession), and column 3-4 is phenotypic data. Data was obtained from Angelovici et al. (2013).

Example file (05_22_2019_Arabidopsis_360_BCAA_raw.csv from tutorial data set):

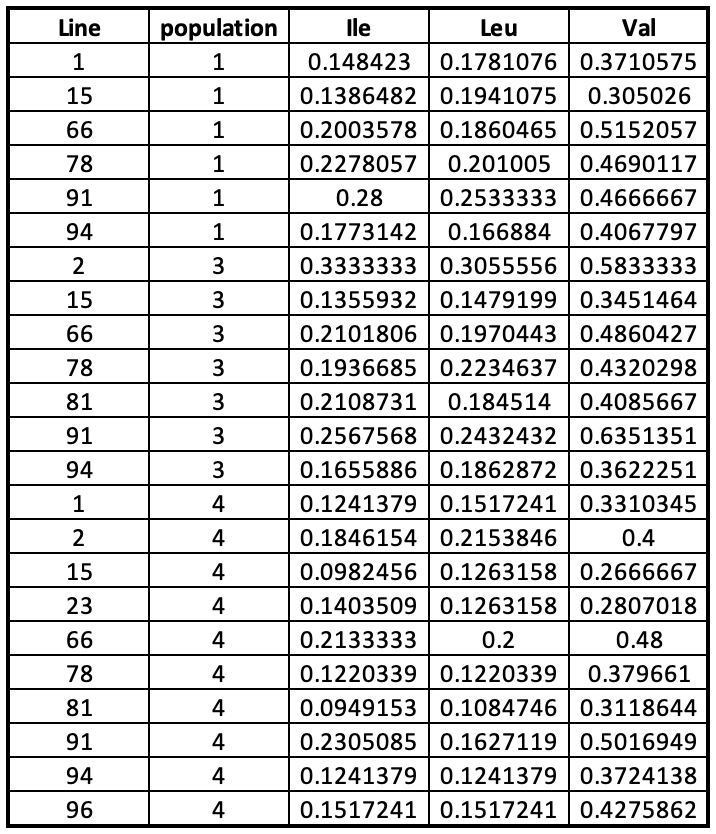

*BLUP/BLUE Data*

1. First column: Accession ID/Taxa name should be the first column of the phenotype file.
2. Subsequent columns: should contain the observed phenotype measurements from each accession where each column designates one trait. Column names will indicate phenotype names to be used in the remaining analysis.

An example of maize BLUP data is provided below where column 1 is the Taxa name and column 2-4 is phenotypic data. Maize data was obtained from Flint-Garcia et al. (2005).

Example file (mdp_traits.txt from tutorial data set):

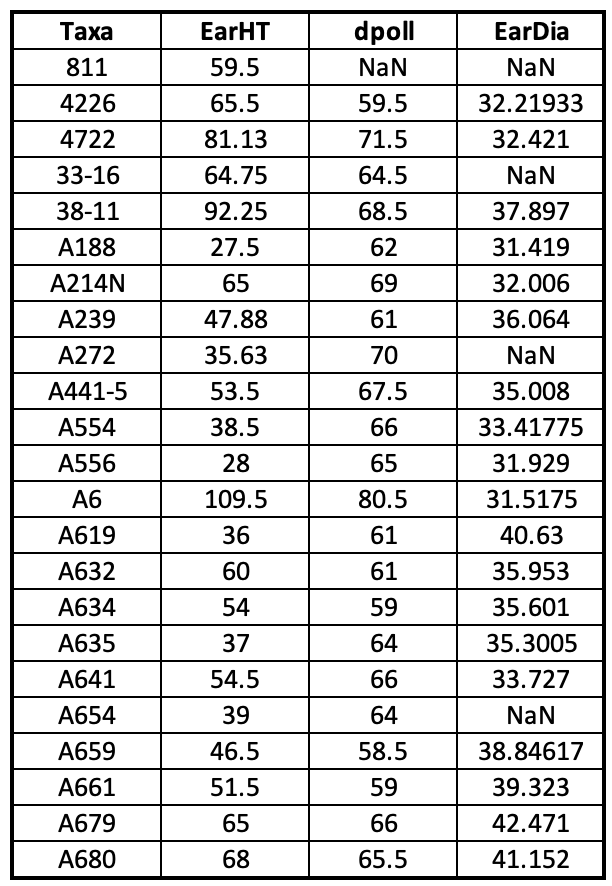

2.2. ***Genotypic data***

Formatted genotypic data from the Goodman Buckler maize association population (Flint-Garcia et al., 2005) are provided with the HAPPI GWAS download. Formatted genotypic data from the *Arabidopsis* 360 population (Horton et al., 2012; Nordborg et al,. 2005) and *Arabidopsis* 1001 population (Alonso-Blanco et al., 2016) is available for download at the Angelovici CyVerse account found here: /iplant/home/angelovici_lab/HAPPI_GWAS

The user can also provide private genotypic data. GAPIT accepts genotypic data in the HapMap or numeric format.

2.2.1. *Hapmap format*

Hapmap format is commonly used to store genotypic data. The first 11 columns of the file are SNP attributes, and the remaining columns are SNP information. The first four columns are SNP identifier, SNP alleles, chromosome number, and position number. These attributes cannot have empty values. The remaining columns can have empty values if the values are not identified. The SNP information in the HapMap consists of genotyped SNPs’ alphabetical codes for the sampled population. Those codes can be encoded using either single-bit or double-bit of the standard IUPAC code.

An example of HapMap format from the maize genotypic data is as follows (mdp_genotype_chr1.hmp.txt):

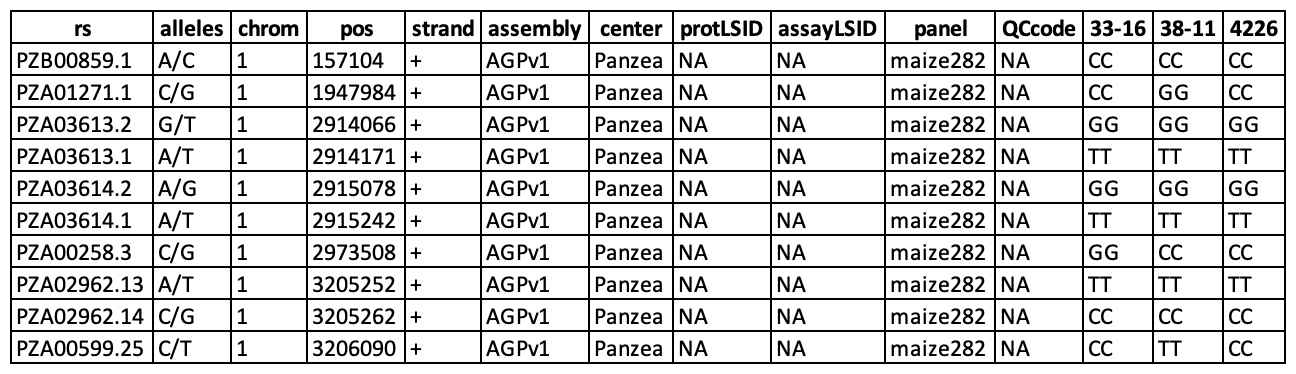

2.2.2. *Numeric format*

In numeric format, columns represent SNPs, and rows represent Accession ID. If numeric genotype data is used, two separate files must be input. The first file contains numeric genotypic data (referred to as the “GD” file). Homozygotes are denoted by “0” and “2” and heterozygotes are denoted by “1”. The second file (referred to as the “GM” file) contains information regarding SNP positions in the genome. The SNP order must be the same in the two files.

An example GD file from the *Arabidopsis* 360 genotypic data is as follows (Call_Method_75_GD1.txt):

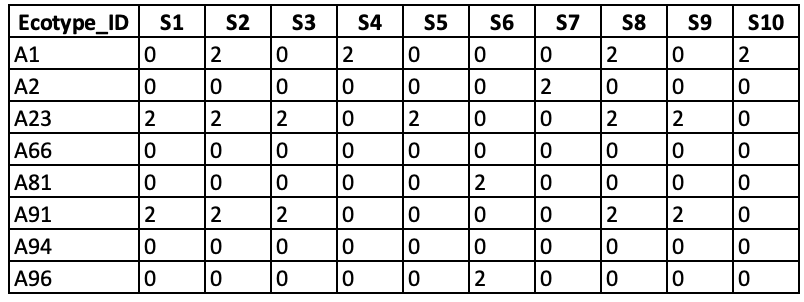

An example GM from the *Arabidopsis* 360 genotypic data is as follows (IMPORTANT_Call_Method_75_GM1.txt):

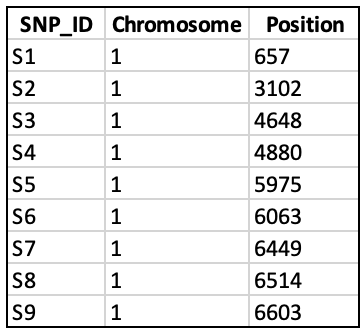

2.2.3. *Naming and importing large genotypic data*

Genotype files that are too large to import due to memory limitations, can be saved per chromosome. Files must be named sequentially (EX: XXX_chr1.hmp.txt, XXXX_chr2.hmp.txt, etc), where the common filename is XXX, and the filename extension (.hmp.txt) are passed to GAPIT using the YAML file. Example genotype files naming are given for the maize (mdp_genotype_chr[1-10].hmp.txt) and *Arabidopsis* 360 populations (Chr[1-5].txt).

For numeric format, the common name and extension file name is passed to GAPIT using “file.GD” and “file.Ext.GD”, respectively; for genotype map files, common name and extension file name is passed to GAPIT using “file.GM” and “file.Ext.GM”, respectively.

Note that it is important that Accession IDs/Taxa names are spelled, punctuated, and capitalized in an identical format in the phenotypic data files as in the genotypic data files. If IDs between the two files do not match, they will be excluded from the analysis.

3. **ANALYSIS**
   3.1. ***Pre-GWAS Analysis***
        3.1.1. *Outlier removal*

Population-wide outlier removal is automatically completed with the *generateBLUP* and *generateBLUE* options. Data are fit using a mixed linear model (see *BLUPs/BLUEs* 3.1.3 for more information on model flexibility) where the influence of data points is determined by using the Studentized deleted residuals (Kutner et al., 2004). That is, a Studentized deleted residual is calculated for every experimental unit. All experimental units where the Studentized residual exceeds the critical value of a null *t*-distribution (for testing$H_{0}$: A given experimental unit is an outlier) with *n-3* degrees of freedom (estimating the intercept, the line variance component, and the population variance component) at a Bonferroni-corrected experimental-wise type I error rate of 0.05 are declared to be an outlier and are removed from the data set.

Output phenotype files contain data with outliers replaced with “NA”; in addition, a data file with a list of removed outlier points and a file with outlier residuals can be retrieved in the “generateBLUP” directory created in the output folder.

3.1.2. *Box-Cox Transformation*

Box-Cox transformation is used to transform each trait to meet normality assumptions. A lambda is calculated per trait and used to transform each trait. The model used to calculate BLUPs/BLUEs will be used at this step (see *BLUPs/BLUEs* 3.1.3 for more information).

The *powerTransform* function is part of the *car* package (Fox et al., 2012) and works by using the maximum likelihood-like approach of Box and Cox (Box & Cox, 1964) to select a transformation. Within the function, the options for *Family* should be set to “bcPower” and *Lambdas* set to -2 to 2. The output phenotype file contains transformed phenotypic data. A list of lambdas for each trait in addition to a data file with the transformed data can be retrieved in the “generateBLUP” directory created in the output folder. Box-Cox transformation is automatically run with the *generateBLUP* and *generateBLUE* options. If errors arise during the transformation step, these traits are flagged and the name of the trait is output to “Traits_not_transform.txt”; flagged traits remain in the analysis but are not transformed and continue to BLUPs/BLUEs calculations. Transformations, such as log transformations, can be applied externally and data can be fed into HAPPI GWAS at the GAPIT step.

3.1.3. *BLUPs/BLUEs*

After outlier removal and transformation, genetic values (either as random or fixed effects) are estimated using the *generateBLUP* and *generateBLUE* options, respectively. A general mixed linear model combines information from all relatives measured to improve estimates (see equation below). In doing so, replicates per accession within a given trait are eliminated, and only one value per accession per trait remains.

$$y = x\beta+ Z\mu+ e$$

Where

$y$ = vector of observation (phenotypes)

$x$ = matrix of fixed effects

$\beta$ = vector of fixed effects to be estimated (i.e. year, location, treatment effects)

$Z$ = matrix of random effects

$\mu$ = vector of random effects to be estimated (genetic values)

$e$ = vector of residual errors

A histogram showing the distribution of the BLUPs/BLUE of each trait is automatically created in the output folder under “BLUP_histogram.png” (see *Intermediate output files* *4.1* for example output). Traits that fail to converge during BLUP/BLUE calculations will be listed in the “Traits_not_converge.txt” file in the output folder.

The user can bypass outlier removal and transformation steps and input externally calculated BLUPs/BLUEs (see *Tutorials* 5.1.2 for more information).

3.1.3.1. *BLUPs (Best Linear Unbiased Prediction)*

A linear mixed model is used to predict random effects (û). In BLUP calculations, the Accession ID/Taxa name will be considered a random effect. All additional variables in the model are random. A file containing the BLUP data can be found in the “generateBLUP” directory created in the output folder.

BLUPs can be run by the following:

Rscript HAPPI_GWAS.R -generateBLUP Arabidopsis_360.yaml

3.1.3.2. *BLUEs (Best Linear Unbiased Estimates)*

A linear mixed model is used to estimate fixed effects (β-hat). In BLUE calculations, the Accession ID/Taxa name will be considered a fixed effect. All additional variables are random. A file containing the BLUE data can be found in the “generateBLUP” directory created in the output folder.

BLUEs can be run by the following:

Rscript HAPPI_GWAS.R -generateBLUE Arabidopsis_360.yaml

3.2. ***GWAS***

Customized YAML files can be created to run the different models in GWAS by calling GAPIT3 (Wang & Zhang, 2018). Model type can be selected in the YAML file by simply removing a “#” in front of the desired and adding a “#” in front of the undesired model(s) (see *Tutorials* 5.1.5 for more details).

To run GWAS, use the following:

Rscript HAPPI_GWAS.R -GAPIT -extractHaplotype -searchGenes Arabidopsis_360.yaml

The *GAPIT* option is required to run GWAS. The *extractHaplotype* and *searchGenes* options are optional.

3.2.1. *Models*

1. Generalized Linear Model (GLM): model including only fixed effects. Population structure is defined (Q matrix). Both the marker and population structure are defined as fixed effects in the model. No random effects are found in the model.
2. Mixed Linear Model (MLM): model including both fixed and random effects. Relatedness is conveyed through a kinship matrix (K) as a random effect and population structure (Q matrix) is accounted for as fixed effect using STRUCTURE (Pritchard et al, 2000) or PCA.
3. Multiple Locus Mixed Linear Model (MLMM): model including forward-backward stepwise linear mixed-model to estimate variance components (Segura et al, 2012).
4. Settlement of MLM Under Progressively Exclusive Relationship (SUPER): a model that extracts a small subset of SNPs and uses them in FaST-LMM (Wang et al, 2014).
5. Farm-CPU (FarmCPU): a model using pseudo QTNs is used to iterate between fixed and random effect models (Liu et al, 2016).
   3.2.2. *Additional GWAS Options in GAPIT*

Gallery of GWAS input parameters in GAPIT

| **Parameter** | **Default** | **Option** | **Description** |
| --- | --- | --- | --- |
| GAPIT_kinship_matrix | NULL | User | Kinship matrix |
| GAPIT_covariates | NULL | User | Covariate Variables |
| GAPIT_hapmap | NULL | User | Genotype data in Hapmap format |
| GAPIT_genotype_data_numeric | NULL | User | Genotype data in numeric format |
| GAPIT_genotype_map_numeric | NULL | User | Genotype Map file in Hapmap format |
| GAPIT_hapmap_file_extension | hmp.txt | User | File extension for Hapmap file |
| GAPIT_genotype_data_numeric_file_extension | NULL | User | File extension for genotype data in numeric format |
| GAPIT_genotype_map_numeric_file_extension | NULL | User | File extension for genotype data in Hapmap format |
| GAPIT_hapmap_filename | NULL | User | File name for genotype data in Hapmap format |
| GAPIT_genotype_data_numeric_filename | NULL | User | File name for genotype data in numeric format |
| GAPIT_genotype_file_path | ./Demo/ | User | Path to genotype file |
| GAPIT_genotype_file_named_sequentially_from | 1 | User | Starting number of sequentially named genotype files |
| GAPIT_genotype_file_named_sequentially_to | 10 | User | Ending number of sequentially named genotype files |
| GAPIT_model | MLM | GLM  MLM  MLMM  SUPER  FarmCPU | GWAS model |
| GAPIT_SNP_MAF | 0.05 | >0 and <1 | Minor allele frequency to filter SNPs |
| GAPIT_PCA_total | 0 | >0 | Number of PC’s as covariates |
| GAPIT_Model_selection | TRUE | TRUE/FALSE | Forward model selection is done using Bayesian information criterion (BIC) to determine optimal PC/Covariables. |
| GAPIT_SNP_test | TRUE | TRUE/FALSE | Perform SNP testing |
| GAPIT_file_output | TRUE | TRUE/FALSE | Provides automatic GAPIT output files |
| GAPIT_p_value_threshold | NULL | >0 and <1 | P-*value* threshold used to filter significant SNPs |
| GAPIT_p_value_fdr_threshold | 0.05 | >0 and <1 | FDR threshold used to filter significant SNPs |
| GAPIT_LD_number | 100000 | >1 | Range (in bp) around significant SNP for LD analysis |

3.3. ***Post-GWAS***
     3.3.1. *Haploblock analysis*

When interpreting significant SNP-trait associations from GWAS, it is beneficial to focus beyond the identified SNP and determine the extent of linkage disequilibrium (LD) surrounding the SNP. SNPs (and genes)(see *Identify Genes 3.3.2* below) contained within this region of high LD are all of the putative interests. When the *extractHaplotype* option is used, for each significant SNP identified in GWAS, pairwise LD is calculated between the significant SNP and every neighboring SNP in a user-defined window using Haploview (Barrett et al, 2004). Regions of high LD (95% confidence bounds on D prime) (i.e. haploblocks) are identified and automatically used downstream in the *searchGenes* section. SNPs are filtered at a 5% minor allele frequency (MAF) and LD is calculated using D prime. Genes contained or partially contained within haploblocks are identified and output with respective gene descriptions in the final summary datasheet. If no genes overlap the haploblock or the significant SNP does not fall within a haploblock, the gene directly upstream and downstream of the significant SNP is given. If no gene annotation file is available, haploblock analysis and gene identification steps can be skipped entirely.

In species with limited genomic information available that prevents accurate LD calculations, the haploblock analysis can be skipped (by removing the *extractHaplotype* option, while the *searchGenes* option is still used) and genes contained in a user-defined window, flanking the significant SNP, can be output. The command is as follows:

Rscript HAPPI_GWAS.R -generateBLUP -GAPIT -searchGenes Arabidopsis_360.yaml

The haploblock section requires a reference file encoded in numerical format. The file is a simpler version of the HapMap format. It only contains two important attributes which are the chromosome number and position number. These two attributes can be found in the first two columns. The remaining columns are samples with their respective SNP information encoded in numerical format. User-defined VCF files can also be used and configured in the YAML file. See *Tutorials* 5.1 and 6.3 for editing YAML files and formatting user input VCF file.

An example file for haploblock analysis from the maize data is as follows (mdp_genotype_haploview_chr1.txt):

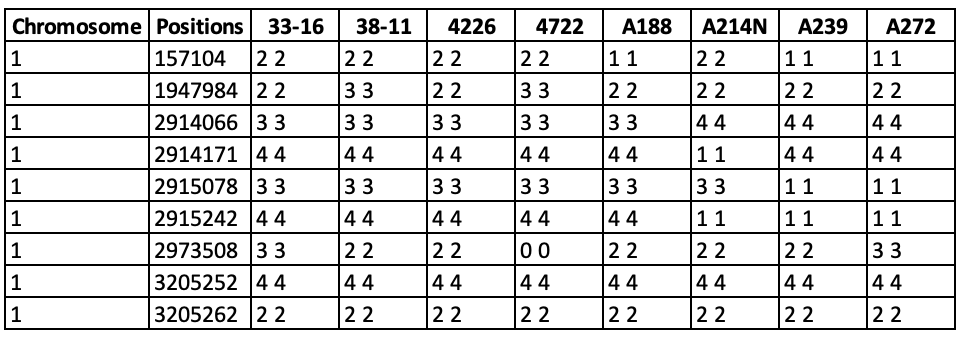

For additional information regarding Haploview, please refer to the official Haploview manual: <https://www.broadinstitute.org/haploview/user-manual>

Gallery of Haploview input parameters

| **Parameter** | **Default** | **Option** | **Description** |
| --- | --- | --- | --- |
| Haploview_file_path | ./Demo/ | User | Path to haploview file |
| Haploview_file_name | mdp_genotype_haploview_chr | User | File name for haploview file |
| Haploview_file_extension | txt | User | File extension for Haploview file |
| Haploview_file_named_sequentially_from | 1 | User | Starting number of sequentially named Haploview files |
| Haploview_file_named_sequentially_to | 10 | User | Ending number of sequentially named Haploview files |

3.3.2. *Identify genes*

Haploblock information for each SNP is automatically used in the *searchGene* option where genes contained in or overlapping with the calculated haploblock (from the *extractHaplotype* section) are identified. By identifying each gene associated with the GWAS significant SNPs, HAPPI GWAS outputs a list of genes, rather than SNPs, which is more informative in determining the complex SNP-trait relationships. Files required to run Haploview in each of the Demo datasets are provided.

LD parameters can be altered through the *GAPIT_LD_number* option in the YAML file. At the *Identify Genes* step, a GFF file is also required. *M*aize GFF files are included in the tool packages. *Arabidopsis* GFF files can be downloaded from Cyverse (See 5.2.2). User-defined GFF files can also be used and designated in the YAML file. See *Tutorials* 5.1 and 6.2 for editing YAML files and formatting user input GFF files.

An example GFF from the maize data is as follows (gene_chr1.gff.txt):

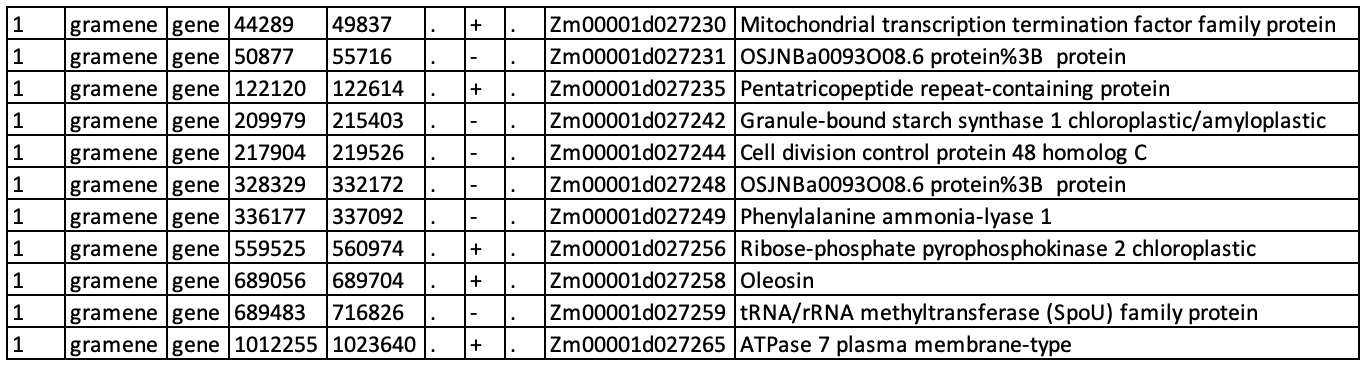

Gallery of GFF file parameters

| **Parameter** | **Default** | **Option** | **Description** |
| --- | --- | --- | --- |
| GFF_file_path | ./Demo/ | User | Path to GFF file |
| GFF_file_name | gene_chr | User | File name for GFF file |
| GFF_file_extension | gff.txt |  | File extension for GFF file |
| GFF_file_named_sequentially_from | 1 | User | Starting number of sequentially named GFF files |
| GFF_file_named_sequentially_to | 10 | User | Ending number of sequentially names GFF files |

4. **RESULTS**

An output directory is automatically generated based on the path provided in the “Output Directory” line of the YAML file. The output directory can be named by the user. The name “demo_output_GLM” (nested in the “output” folder) is used in the maize demo YAML file. When the tool is running, some folders and subfolders are generated within the output directory to store results and temporary files. In this section, only the folders and subfolders used to store the final output will be discussed here.

4.1. ***Intermediate output files***

The folder “generateBLUP” found within the automatically created output folder contains intermediate files that are created throughout the pre-GWAS portion of the pipeline. These files include:

1. A text file containing outlier residuals produced from Studentized Deleted Residuals analysis (Outliers_residuals.txt).
2. A data file containing phenotype data with outliers removed (Outlier_removed_data.csv).
3. A list of data points identified as outliers through Studentized Deleted Residuals analysis (Outlier_data.csv).
4. A list of lambda values calculated for each trait during Box-Cox transformation (Lambda_values.csv).
5. A data file containing Box-Cox transformed phenotype data (Boxcox_transformed_data.csv).
6. A data file containing a list of traits that failed to transform during Box-Cox transformation (Traits_not_transform.txt).
7. A data file containing calculated BLUPs (if *generateBLUP* flag is used) (BLUP.csv).
8. A data file containing calculate BLUEs (if *generateBLUE* flag is used) (BLUE.csv).
9. A data file containing a list of traits that failed to converge during BLUP/BLUE calculations (Traits_not_converge.txt).
10. A png file containing histograms of the BLUP/BLUE distributions for each trait (BLUP_histogram.png/BLUE_histogram.png).

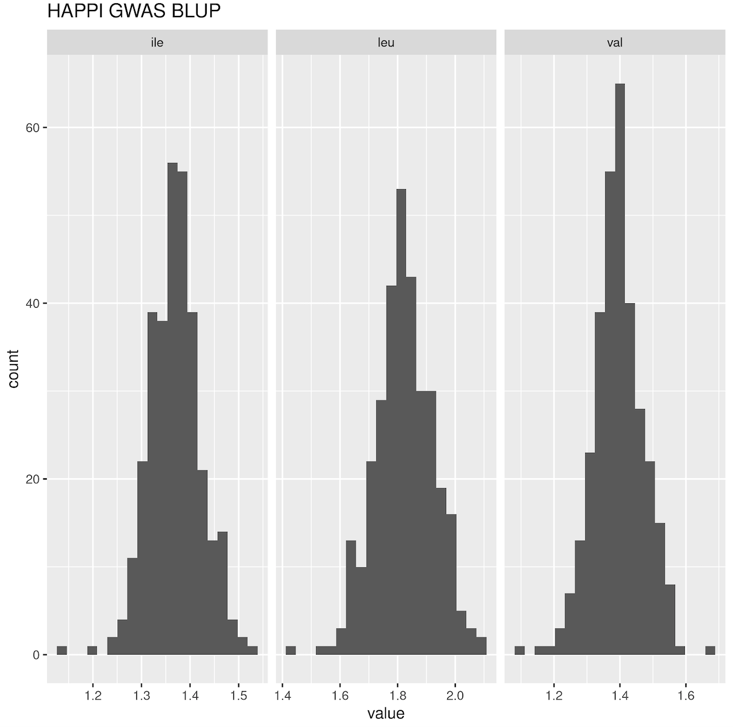

**Figure 4.1-1:** Summary histograms showing the BLUP/BLUE distribution for each trait.

4.2. ***GWAS output***

The subfolder “GAPIT” under the main output folder contains all data resulting from the GWAS and post-GWAS analysis. HAPPI GWAS creates publication-ready summaries and figures, in addition to providing GAPIT output. Only unique HAPPI GWAS files are described here. For more information on GAPIT output figures and tables, please refer to the GAPIT manual (Wang & Zhang, 2018). HAPPI GWAS output files using the maize demo data are shown below:

4.2.1. HAPPI GWAS summary figures and tables

1. GAPIT.combined.GWAS.Results.csv

**Table 4.2.1-1:** Summary output combining GWAS and haploblock results across all traits. All significant SNPs identified at the user-defined threshold with their associated traits, genes, and gene descriptions are provided.

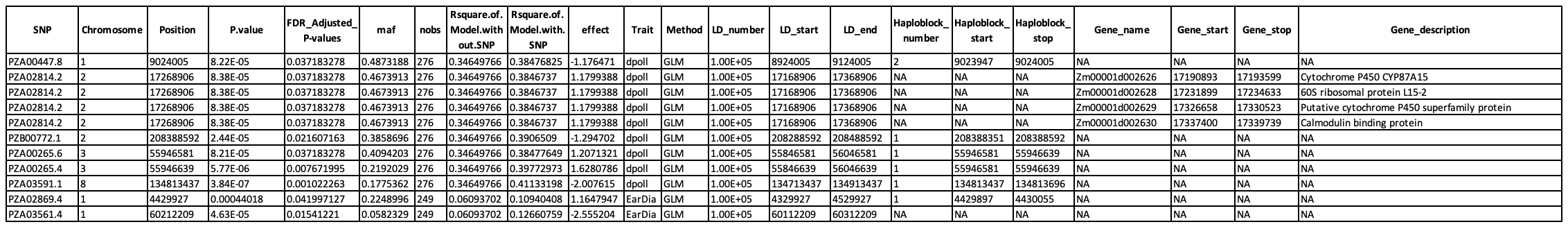

2. Summary Heatmap

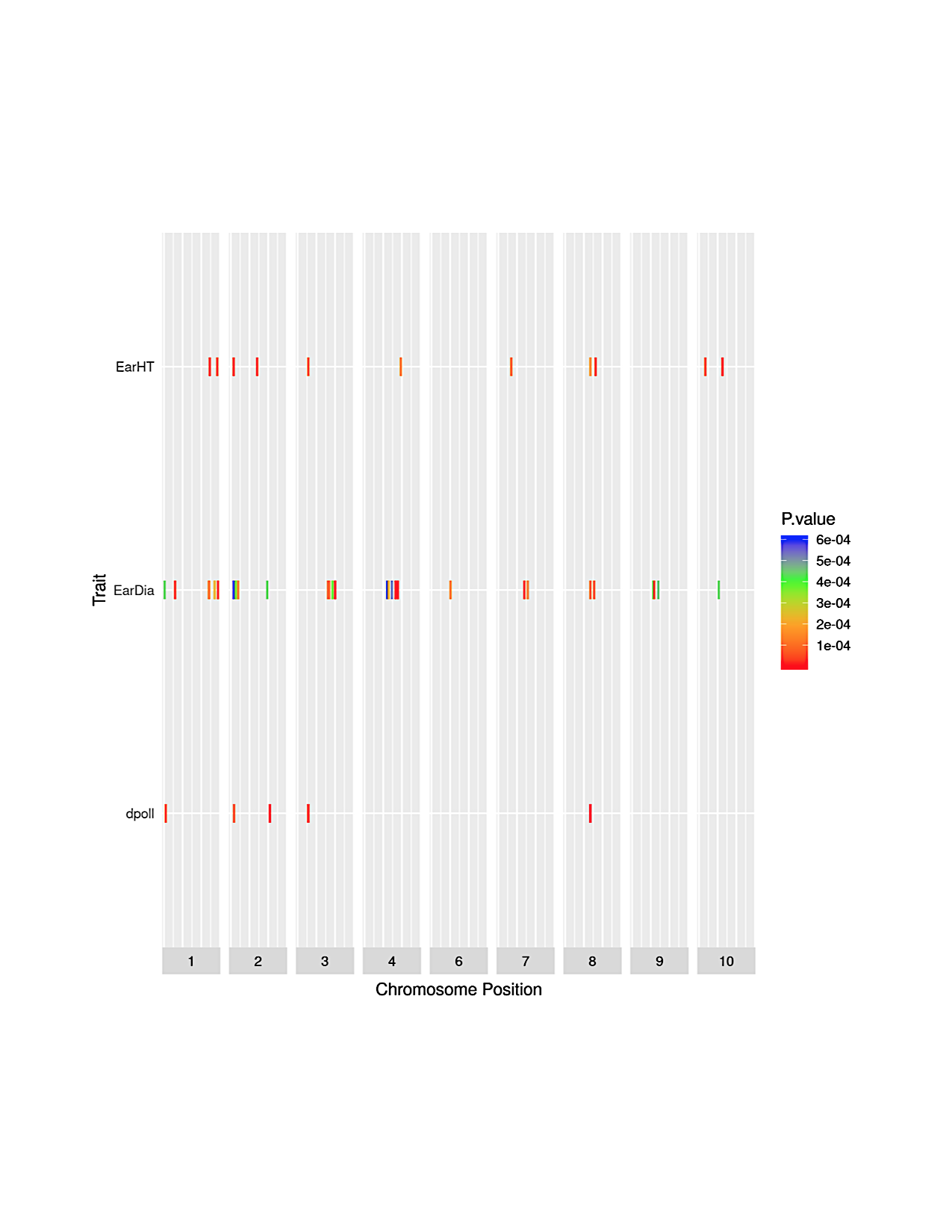

**Figure 4.2.1-1:** Summary heatmap illustrating significant SNP distribution across chromosomes from all traits. The x-axis shows only chromosomes containing significant SNPs; the y-axis shows all traits with significant SNP-trait associations. Rectangles represent SNPs that are color-coded based on P-*value*.

3. Most frequently occurring summary table

**Table 4.2.1-2:** Summary output including GWAS and haploblock results for the top five most recurring significant SNPs across all traits.

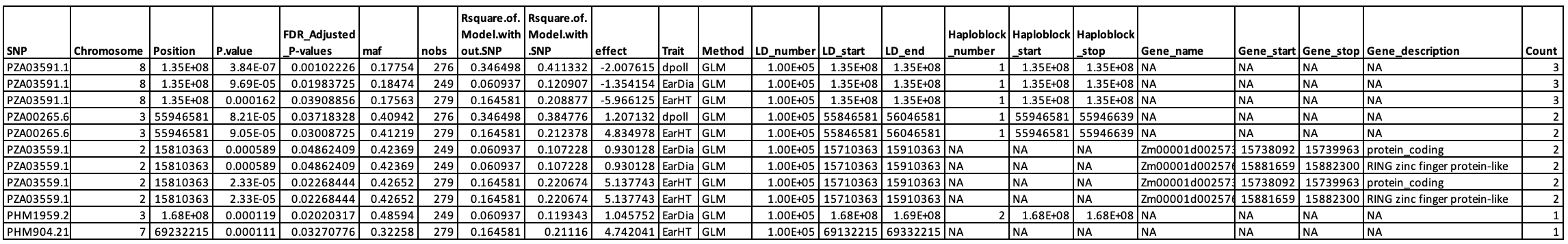

4. Lowest p-value summary table

**Table 4.2.1-3:** Summary output including GWAS and haploblock results for the top five significant SNPs across all traits with the lowest P-*value*.

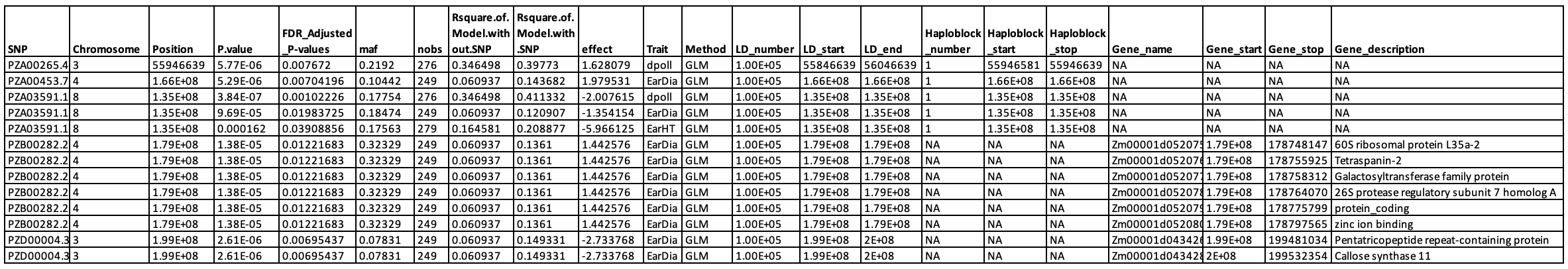

4.2.2. *Additional output*
       4.2.2.1. Haploview_Haplotypes_gabriel_blocks

This folder contains text files that have haplotype block information. The haploblock information includes the total number of haploblocks and markers for each significant SNP.

4.2.2.2. Haplotview_LD_plot

This folder contains LD plots. The LD plots are the visualizations of the haploblocks for each significant SNP.

4.2.2.3. GAPIT_auto_output

GAPIT plots, figures, and tables for each trait.

5. **TUTORIALS**
   5.1. ***Editing the YAML file***

The YAML file is a configuration file that includes the names and locations of all datafiles and controls data input and output at each step. It must be stored directly inside the yaml folder which is under the main HAPPI_GWAS directory. The YAML file is composed of six sections: Raw Data, BLUP or BLUE, GAPIT3, Haploview, Match Gene Start and Stop, and Output Directory.

5.1.1. *Inputting raw data*

If starting with raw data, the “Raw Data” section of the YAML file must be edited so that the “raw_data” line includes the path and file name of the raw data. The line “by_column” designates any descriptive columns that are not phenotypic data. “start_column” designates the first column with phenotypic measurements.

In the example below, columns 1 and 2 correspond to Accession ID and replicate, respectively; column 3 marks the first trait with subsequent traits in following columns.

When raw data is provided, there should be no BLUP/BLUE file given. Note in the example, under the “BLUP or BLUE” section, the line “BLUP” is blank.

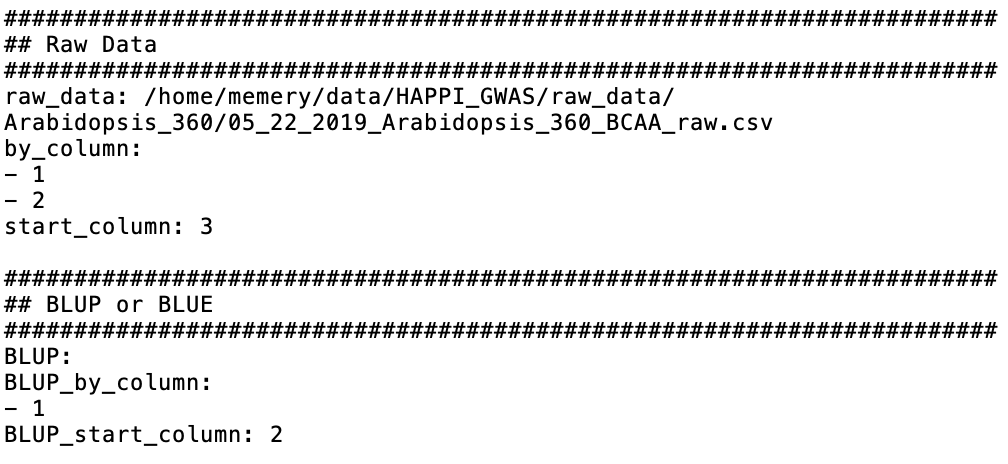

5.1.2. *Inputting BLUP/BLUE data*

Instead of beginning with raw data, externally calculated BLUPs/BLUEs can be directly imported into the pipeline, bypassing upstream analyses required for raw data. To bypass importing raw data, keep “raw_data” blank in the “Raw Data” section. In the “BLUP or BLUE” section, add the path to the externally calculated BLUPs/BLUEs. In the BLUP/BLUE file, the first column should be the Accession ID, and subsequent columns should contain the BLUPs/BLUEs. The “BLUP” line accepts both BLUP and BLUE calculations. An example outputting BLUP calculations is shown below.

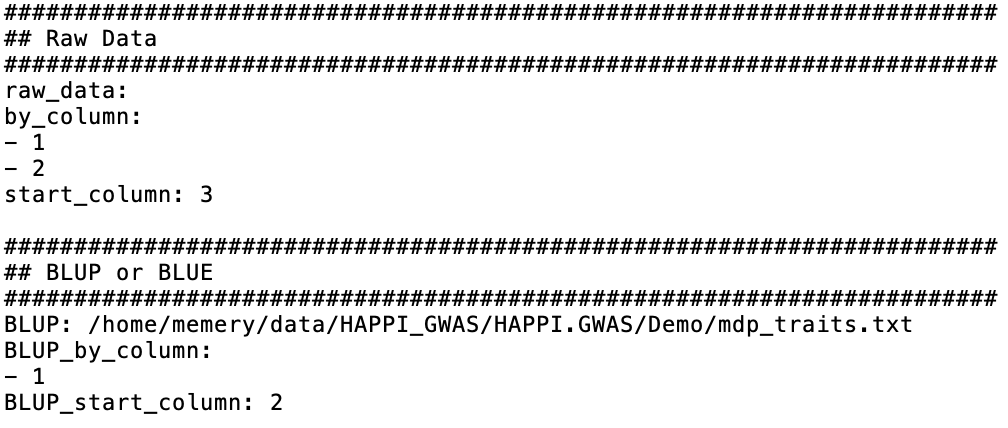

5.1.3. *Editing paths in the YAML file*

There are six instances where the file path needs to be edited by the user in the YAML file. Failure to change one path will result in the pipeline not running to entirety. Below is a list of sections and lines that need to be edited:

Gallery of paths to be edited in the YAML file

| **YAML section** | **Line in YAML** | **Notes** |
| --- | --- | --- |
| Raw Data | raw_data | Required if starting with raw data. Should be blank if BLUPs/BLUEs are provided. |
| BLUP or BLUE | BLUP | Required if starting with BLUPs/BLUEs data and bypassing upstream analysis. Should be blank if raw data are provided. |
| Gapit3 | GAPIT_genotype_file_path | Path to genotype file |
| Haploview | Haploview_file_path | Path to Haploview file |
| Match Gene Start and Stop | GFF_file_path | Path to GFF file |
| Output Directory | output | Path where output directory will be created and all output will be stored. |

5.1.4. *Filtering by P*-*value or FDR*

HAPPI GWAS has the capacity to filter significant SNPs based on FDR or P-*value*. In the YAML file under the “Gapit3” section, a user-defined P-*value* threshold can be added at “GAPIT_p_value_threshold” while “GAPIT_p_value_fdr_threshold” is left blank. If a FDR cutoff is desired, a user-defined FDR threshold can be added at “GAPIT_p_value_fdr_threshold” while “GAPIT_p_value_threshold” is left blank. To filter by a Bonferroni cutoff, simply take the desired P-*value* threshold and divide it by the total number of SNPs. Put this calculated number (Bonferroni corrected P-*value* threshold) under “GAPIT_p_value_threshold”.

5.1.5. *Editing the model*
       5.1.5.1. *Number of variables in the model*

Additional variables can be added to the model in the “Raw Data” or “BLUP or BLUE” section. By designating the column number where phenotypic values begin, HAPPI GWAS will automatically assume all prior columns (except column 1 that contains Accession ID) are variables in the model.

Below is an example showing the “Raw Data” section in a YAML file with phenotype data starting in column 5; column 1 - 4 will be used as variables in the model in the BLUP/BLUE model. If the *generateBLUP* option is used, column 1 will be considered a random effect. Conversely, if the *generateBLUE* option is used, column 1 will be considered a fixed effect. Columns 2 - 4 will be considered a random effect regardless if the *generateBLUP* or *generateBLUE* option is used.

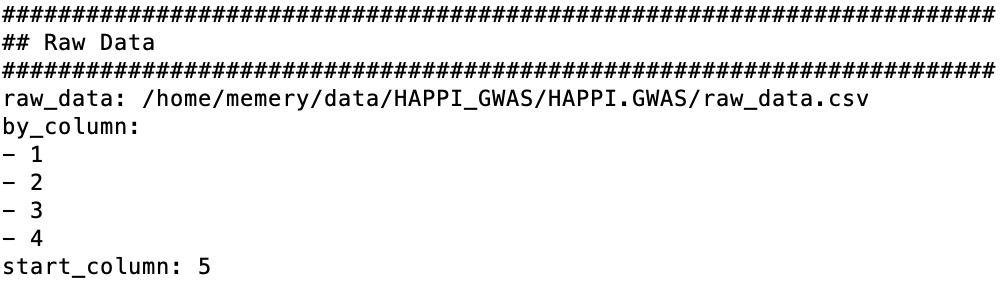

Below is an example showing the “BLUP or BLUE” section in a YAML file depicting phenotype data starting in column 2; column 1 will be used as a variable in the model of subsequent steps.

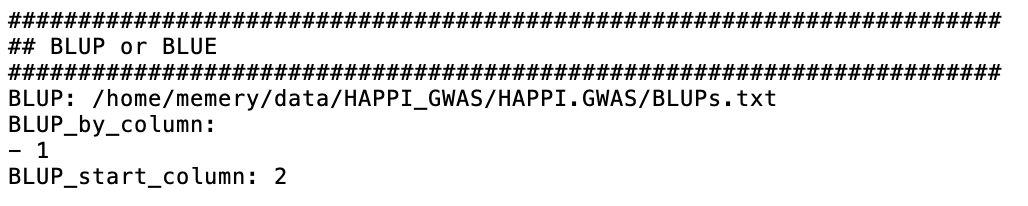

5.1.5.2. *Fixed versus random variables in the model*

In the GWAS analysis, each step in pre-GWAS should fit the same model; these steps include outlier removal, Box-Cox transformation, and BLUP/BLUE calculations. In BLUP calculations, line/accession is considered a random effect in the model. Contrastly, in BLUE calculations, line/accession is considered a fixed effect in the model. HAPPI GWAS automatically chooses the correct model at each pre-GWAS step based on user-defined parameters identified at the raw data input step and/or through the usage of the *generateBLUP* or *generateBLUE* options.

5.2. ***Tutorial data sets***

All necessary inputs to run the maize demo data are provided within the cloned repository from GitHub. Input to run the *Arabidopsis* demo data can be downloaded from Cyverse (See 5.2.2). Files include phenotype data, genotype data, Haploview files, and GFF gene annotation files.

5.2.1. *Demo maize*

Maize Demo files can be found in the *Demo* folder within the cloned HAPPI GWAS GitHub repository. Hapmap, Haploview, and GFF files are provided and split by chromosome into 10 files, respectively. BLUPs have been externally calculated. All phenotypic data was obtained from Flint-Garcia et al. (2005). For this tutorial, navigate into the cloned HAPPI GWAS repository (refer to *Getting Started* 1.2 to determine the absolute path). Please refer to the following commands:

```
#navigate into the user-created HAPPI_GWAS folder
cd <absolute path identified before the HAPPI.GWAS repository cloning process>
cd HAPPI.GWAS	# navigate into the cloned repository
```

To run the Maize Demo data follow these steps:

Step 1: Edit the Demo_GLM.yaml file:

a. Edit the “BLUP or BLUE” section. Ensure the path and file name are correct.

In this tutorial, we will be using externally calculated BLUPs (i.e. the “Raw Data” section is blank). The first column in the BLUPs file is the Line ID, and subsequent columns (starting with column 2) are the phenotypic data in the form of BLUPs.

b. Edit the “GAPIT3” section. Ensure the path at line “GAPIT_genotype_file_path” is correct (i.e. using the correct absolute path). We will be using mdp_genotype_chr[1-10].hmp.txt files. Note how the *GLM* is the selected model as all other model options are ignored by the addition of #. SNP MAF is filtered at 0.05 with a significant FDR threshold of 0.05. A desired window size of 100,000 bp on each side of the significant SNP is defined by editing GAPIT_LD_number: 100000.
c. Edit the “Haploview” section. Ensure the path at line “Haploview_file_path” is correct. We will be using mdp_genotype_haploview_chr[1-10].txt files.
d. Edit the “Match Gene Start and Stop” section. Ensure the path at line “GFF_file_path” is correct. We will be using gene_chr[1-10].gff.txt files.
e. Edit the “Output Directory” section. Ensure the path on line “output” is correct.

Below is the example Demo.GLM.yaml file.

*Note in the example Demo YAML file below, the absolute path does not require editing. The directory structure of the HAPPI GWAS package is set, and thus paths to the Demo directory are provided for you.*

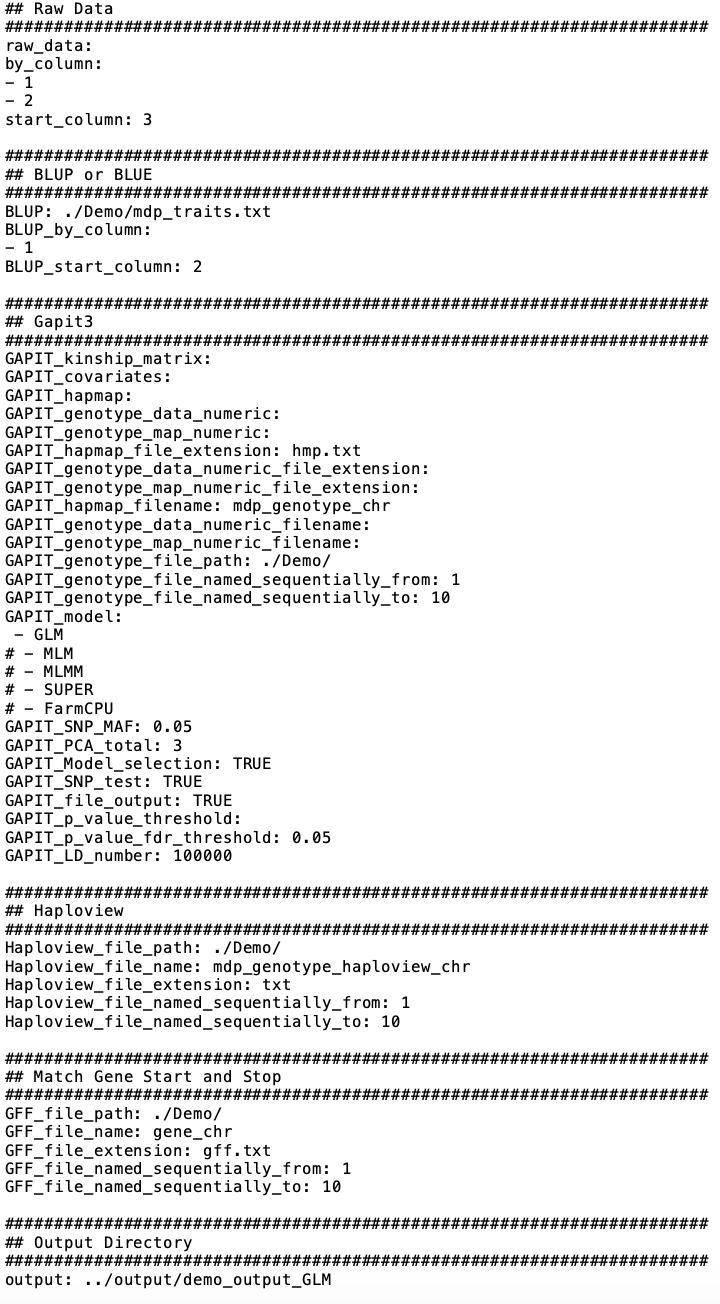

Step 2: Run HAPPI GWAS using the following command:

Rscript HAPPI_GWAS.R -GAPIT -extractHaplotype -searchGenes Demo_GLM.yaml

*Note how the generateBLUP option is not being used here as raw data is not being inputted.*

Step 3: Access output data at the following:

cd <user defined output path found in the “Output Directory” section of the YAML file>

5.2.2. *Demo Arabidopsis 360 population*

To run the *Arabidopsis* 360 Demo, users must download the raw data file, genotype data files, genotype map files, Haploview files, and GFF files from Cyverse (user must first complete *Getting Started 1.2* ). In order to do so, users must first create an account with Cyverse. Copy and paste the following link in the browser and click register to make a new account. Users who already have a Cyverse account can ignore this step.

<https://de.cyverse.org/de>

iCommands software needs to be installed on the machine that is being used to download the data. To install iCommands and initialize iRODS connection, copy and paste the following link in the browser and follow the instructions on the website.

<https://wiki.cyverse.org/wiki/display/DS/Setting+Up+iCommands#SettingUpiCommands-Installation>

After installing the iCommands software and initializing the iRODS connection, users must navigate to the reference files directory and clone the required files (the raw data file, genotyped data files, genotyped map files, Haploview files, and GFF files) from Cyverse. The genotype data, genotype map, Haploview, and GFF files are split by chromosome into five files (for the five chromosomes in the *Arabidopsis* genome). *Arabidopsis* data is provided in raw form. Raw phenotypic data was obtained from Angelovici et al. (2013). Please use the commands below to complete all the download processes.

cd <absolute path noted in Getting Started 1.2>

cd reference_files

icd /iplant/home/angelovici_lab/HAPPI_GWAS/

iget -rfvTK /iplant/home/angelovici_lab/HAPPI_GWAS/gene_annotation_files ./

iget -rfvTK /iplant/home/angelovici_lab/HAPPI_GWAS/genotype_files ./

iget -rfvTK /iplant/home/angelovici_lab/HAPPI_GWAS/haploview_files ./

cd ../raw_data

iget -rfvTK /iplant/home/angelovici_lab/HAPPI_GWAS/raw_data/Arabidopsis_360 ./

cd ../HAPPI.GWAS

After successfully downloading all files, users are now ready to run the *Arabidopsis* 360 Demo data. To run the *Arabidopsis* 360 Demo data follow these steps:

Step 1: Edit the Arabidopsis360.yaml file:

a. Edit the “Raw Data” section. Ensure the path and file name are correct.

In this tutorial, we will be using raw data (i.e. the “BLUP or BLUE” section is blank). The first column in the data file is the *Line* (i.e. Accession ID), the second is the *population* (i.e. replicate), and subsequent columns are the phenotypic data in raw form.

b. Edit the “GAPIT3” section. Ensure the path at line “GAPIT_genotype_file_path” is correct. We will be using genotype data in the numeric format. Files named Call_Method_75_GD[1-5].txt files will be used. Note how the *MLM* is the selected model as all other model options are ignored by the addition of #. SNP MAF is filtered at 0.05 with a significant FDR threshold of 0.05. An average LD decay of 5,000 bp if used; therefore, we chose a GAPIT_LD_number of 5000 (bp on each side of the significant SNP).
c. Edit the “Haploview” section. Ensure the path at line “Haploview_file_path” is correct. We will be using Chr[1-5].haploview.txt files.
d. Edit the “Match Gene Start and Stop” section. Ensure the path at line “GFF_file_path” is correct. We will be using Chr[1-5].txt files.
e. Edit the “Output Directory” section. Ensure the path on line “output” is correct.

Below is the example Arabidopsis360.yaml file.

*Note in the example YAML file below that the absolute path is as follows: /home/memery/data/HAPPI_GWAS/*

*The absolute path will change for every user (See Getting Started 1.2).*

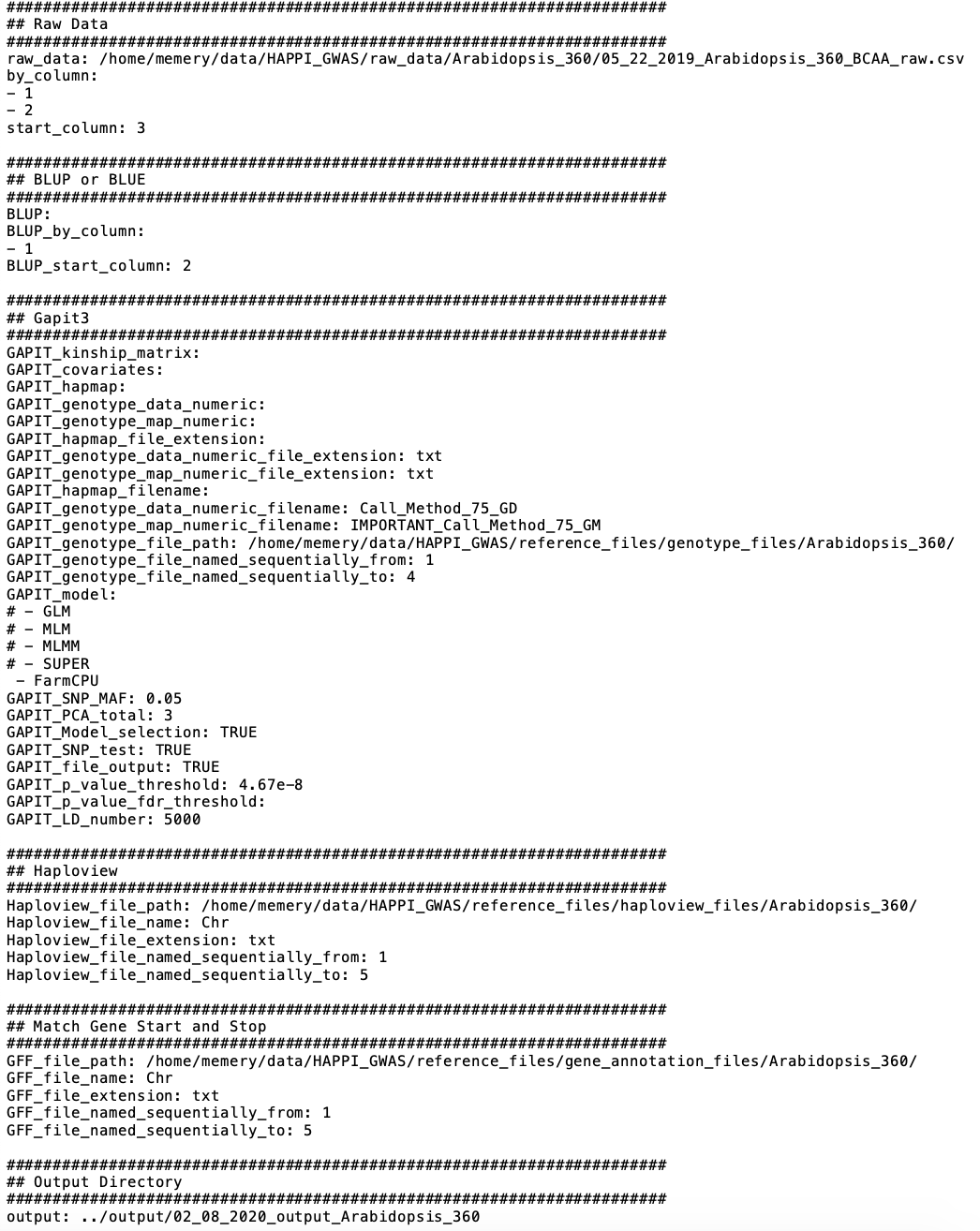

Step 2: Run HAPPI GWAS using the following command (ensure command is run from the HAPPI.GWAS directory):

Rscript HAPPI_GWAS.R -generateBLUP -GAPIT -extractHaplotype -searchGenes Arabidopsis360.yaml

Note how the *generateBLUP* option is used. Therefore, *Line* and *population* are fit as random effects in the model. The “BLUP or BLUE” section is blank.

Step 3: Access output data at the following:

cd < user-defined output path found in the “Output Directory” section of the YAML file >

5.2.3. *Arabidopsis 1001 population*

Formatted genotype data, Haploview files, and GFF gene annotation files for the *Arabidopsis* 1001 population (Alonso-Blanco et al., 2016) can be downloaded from the CyVerse account. Original genotype data was obtained from the 1001 Genomes website (<https://1001genomes.org/data-center.html>) and was filtered at 20% SNP call rate and a MAF of 0.05.

Formatted genotype data, Haploview files, and GFF gene annotation files downloaded and used in the *Arabidopsis* 360 demo (See *Demo Arabidopsis 360 Population* 5.2.2) can also be used to run the *Arabidopsis* 1001 data. Users who have completed the *Arabidopsis* 360 demo can skip this section. Users who did not complete the *Arabidopsis* 360 demo need to download required reference files for *Arabidopsis* using the following commands:

cd <absolute path noted in Getting Started 1.2>

cd reference_files

mkdir genotype_files

mkdir haploview_files

mkdir gene_annotation_files

icd iplant/home/angelovici_lab/HAPPI_GWAS/

cd genotype_files

iget -rfvTK /iplant/home/angelovici_lab/HAPPI_GWAS/genotype_files/Arabidopsis_1001 ./

cd ../haploview_files

iget -rfvTK /iplant/home/angelovici_lab/HAPPI_GWAS/haploview_files/Arabidopsis_1001 ./

cd ../gene_annotation_files

iget -rfvTK \ /iplant/home/angelovici_lab/HAPPI_GWAS/gene_annotation_files/Arabidopsis_1001 ./

cd ../../HAPPI.GWAS

Modify the YAML file to reflect changed paths and input files (See *Tutorials* 5.1 for more information regarding editing YAML files).

6. **APPENDIX**
   6.1. Properties of tutorial files

**Table 6.1-1:** Properties of Maize Demo tutorial files provided with HAPPI GWAS download

| **File Name** | **Date** | **Time** | **Size (KB)** |
| --- | --- | --- | --- |
| gene_chr1.gff.txt | 02.03.2020 | 4:01 PM | 164 |
| gene_chr2.gff.txt | 02.03.2020 | 4:01 PM | 197 |
| gene_chr3.gff.txt | 02.03.2020 | 4:01 PM | 158 |
| gene_chr4.gff.txt | 02.03.2020 | 4:01 PM | 186 |
| gene_chr5.gff.txt | 02.03.2020 | 4:01 PM | 107 |
| gene_chr6.gff.txt | 02.03.2020 | 4:01 PM | 120 |
| gene_chr67.gff.txt | 02.03.2020 | 4:01 PM | 105 |
| gene_chr8.gff.txt | 02.03.2020 | 4:01 PM | 155 |
| gene_chr9.gff.txt | 02.03.2020 | 4:01 PM | 151 |
| gene_chr10.gff.txt | 02.03.2020 | 4:01 PM | 81 |
| mdp_genotype_chr1.hmp.txt | 02.03.2020 | 4:01 PM | 489 |
| mdp_genotype_chr2.hmp.txt | 02.03.2020 | 4:01 PM | 357 |
| mdp_genotype_chr3.hmp.txt | 02.03.2020 | 4:01 PM | 322 |
| mdp_genotype_chr4.hmp.txt | 02.03.2020 | 4:01 PM | 290 |
| mdp_genotype_chr5.hmp.txt | 02.03.2020 | 4:01 PM | 324 |
| mdp_genotype_chr6.hmp.txt | 02.03.2020 | 4:01 PM | 194 |
| mdp_genotype_chr7.hmp.txt | 02.03.2020 | 4:01 PM | 224 |
| mdp_genotype_chr8.hmp.txt | 02.03.2020 | 4:01 PM | 233 |
| mdp_genotype_chr9.hmp.txt | 02.03.2020 | 4:01 PM | 194 |
| mdp_genotype_chr10.hmp.txt | 02.03.2020 | 4:01 PM | 183 |
| mdp_genotype_haploview_chr1.txt | 02.03.2020 | 4:01 PM | 615 |
| mdp_genotype_haploview_chr2.txt | 02.03.2020 | 4:01 PM | 448 |
| mdp_genotype_haploview_chr3.txt | 02.03.2020 | 4:01 PM | 405 |
| mdp_genotype_haploview_chr4.txt | 02.03.2020 | 4:01 PM | 364 |
| mdp_genotype_haploview_chr5.txt | 02.03.2020 | 4:01 PM | 407 |
| mdp_genotype_haploview_chr6.txt | 02.03.2020 | 4:01 PM | 244 |
| mdp_genotype_haploview_chr7.txt | 02.03.2020 | 4:01 PM | 281 |
| mdp_genotype_haploview_chr8.txt | 02.03.2020 | 4:01 PM | 293 |
| mdp_genotype_haploview_chr9.txt | 02.03.2020 | 4:01 PM | 244 |
| mdp_genotype_haploview_chr10.txt | 02.03.2020 | 4:01 PM | 230 |
| mdp_traits.txt | 02.03.2020 | 4:01 PM | 7 |

6.2. ***Formatting user input GFF file***

1. Reorganize last (9th) column to contain only the gene ID
2. Insert a 10th column that contains the gene description
3. Split the GFF file by Chr and name the following (“Chr”, # = number of segmented file): *Chr#.txt*
   6.3. ***Formatting user input Hapmap file***
1. Use vcftools vcf-to-tab function to convert VCF to a tab-delimited file
2. Convert tab-delimited file to HapMap format using TASSEL command-line software
3. Split the HapMap file by chromosome and name the following (“Chr”, # = number of segmented file): *Chr#.hmp.txt*
   6.4. ***Formatting user input Haploview file***
1. Use vcftools vcf-to-tab function to convert VCF to a tab-delimited file
2. Convert tab-delimited file to HapMap format using TASSEL command-line software
3. Remove the reference column and keep only chromosome, position, and all sample columns
4. Convert IUPAC to numeric codes: A/A = 1; C/C = 2; G/G = 3; T/T = 4
5. Split file by chromosome
   6.5. **Frequently asked questions**
1. **My BLUP model won’t converge?**

A: Low variance among the accessions may lead to model convergent issues. If models do not converge when BLUPs are calculated, we recommend fixing a model with Accession IDs as fixed effects and calculating BLUEs.

2. **My files are in the correct directories, but GAPIT can’t find them. Why?**

A: There is most likely an issue with the file paths in the YAML file. Ensure the path in each section of the YAML file is correct. Also, be sure you are reading in either raw data *or* BLUPs/BLUEs, but not both. Ensure the YAML file is under the main directory (HAPPI GWAS).

3. **I am trying to run the Rscript command on the command line but keep getting the “Rscript not found” error. What should I do?**

A: For Linux users, R will be added to the system path by default. If it is not added to the system path, please type export PATH="/usr/bin/R:$PATH" in the terminal. Please be aware that if R is not installed using the admin privilege, the R path will be different, please find out the R path and replace /usr/bin/R with the correct R path.

4. **I am trying to use the Rscript command to run the HAPPI GWAS pipeline and get the following error: “there is no package called ‘argparse’”. What should I do?**

A: The argparse package in R relies on the argparse package in Python3. Therefore, users should install Python3 on their machine if they received this error.

5. **Can I use the tool on a Windows machine?**

A: This tool has not been tested on a Windows machine. To run the tool on a Windows machine, the developer recommends users to use a virtual machine and install CentOS/Ubuntu on that virtual machine. After that, please follow the instructions on the *1.2 Getting Started* section to run this tool on the virtual machine.
