## Supplementary Figure S1 for "HAPPI GWAS: Holistic Analysis with Pre and Post Integration GWAS"

### Slide 1
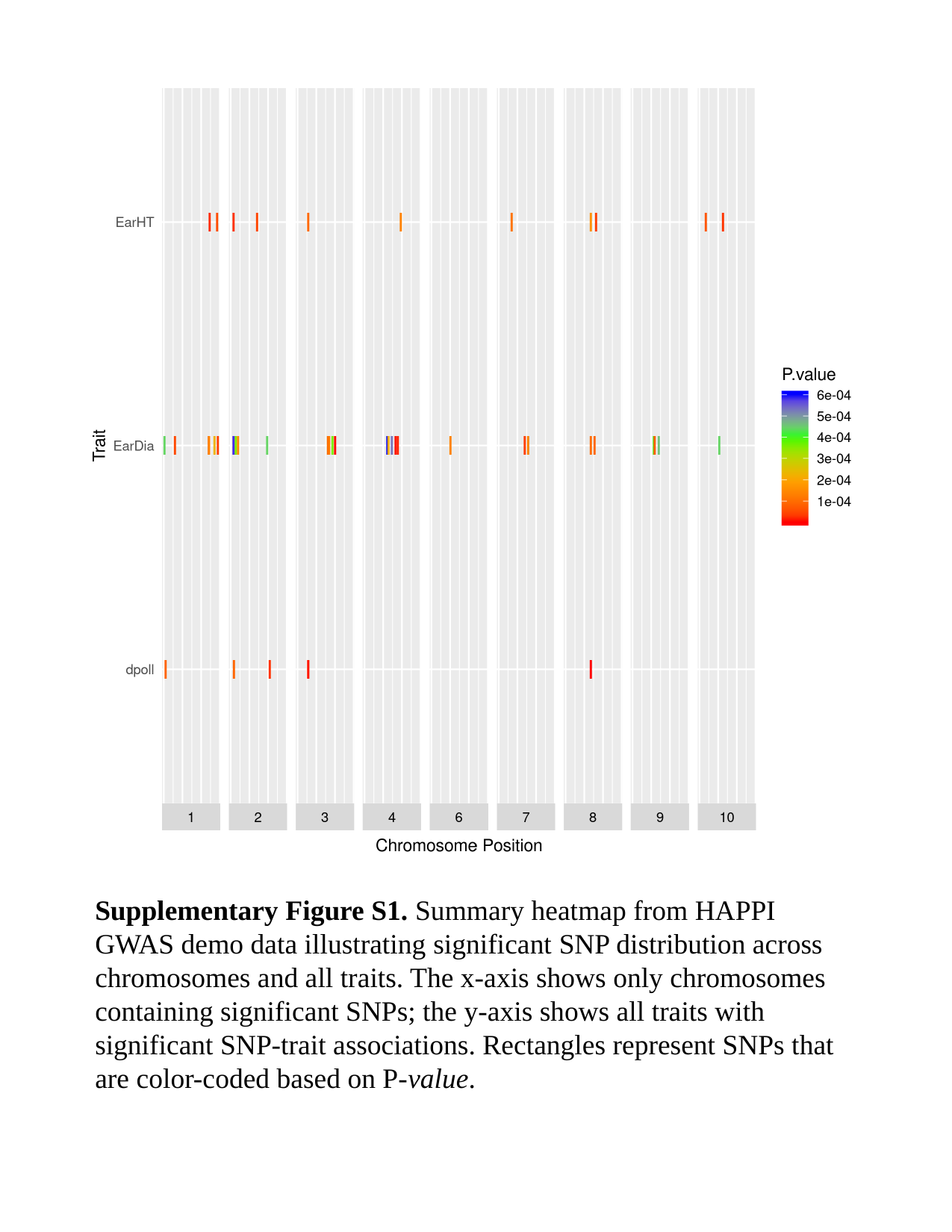

Supplementary Figure S1. Summary heatmap from HAPPI GWAS demo data illustrating significant SNP distribution across chromosomes and all traits. The x-axis shows only chromosomes containing significant SNPs; the y-axis shows all traits with significant SNP-trait associations. Rectangles represent SNPs that are color-coded based on P-value.
